# Real-time search of all bacterial and viral genomic data

**DOI:** 10.1101/234955

**Authors:** Phelim Bradley, Henk C Den Bakker, Eduardo P. C. Rocha, Gil McVean, Zamin Iqbal

## Abstract

Genome sequencing of pathogens is now ubiquitous in microbiology, and the sequence archives are effectively no longer searchable for arbitrary sequences. Furthermore, the exponential increase of these archives is likely to be further spurred by automated diagnostics. To unlock their use for scientific research and real-time surveillance we have combined knowledge about bacterial genetic variation with ideas used in web-search, to build a DNA search engine for microbial data that can grow incrementally. We indexed the complete global corpus of bacterial and viral whole genome sequence data (447,833 genomes), using four orders of magnitude less storage than previous methods. The method allows future scaling to millions of genomes. This renders the global archive accessible to sequence search, which we demonstrate with three applications: ultra-fast search for resistance genes MCR1-3, analysis of host-range for 2827 plasmids, and quantification of the rise of antibiotic resistance prevalence in the sequence archives.

---

Whole genome sequencing (WGS) offers unparalleled resolution for problems as diverse as contact tracing, mapping the spread of drug resistance, identifying zoonoses, and investigating the underlying biology of infectious diseases. Sequence data is deposited in the global sequence archives (European Nucleotide Archive (ENA), Sequence Read Archive (SRA)) which are doubling every two years. This trend is expected to accelerate as affordable WGS-based diagnostic tests become a reality^1-6^. However, it is already impossible to search the archives for datasets with specific mutations (single nucleotide polymorphisms (SNPs)), genes or mobile elements. The ability to combine these queries would be transformative for global management of infectious disease, allowing instant access to datasets within any specified genetic distance (e.g. “has anyone in the world seen something within 20 SNPs of this strain before?), or with given drug resistance mutations, genes or plasmids. A key realization is that this can be achieved by focusing first on exact-match search (“is this precise sequence present?”) rather than alignment.

This formulation, as an exact-search problem, is not a traditional approach in biology. In the 1990s, when most species had one reference genome and within-species variation was less of a focus, BLAST^7^ and its successors^8,9^ revolutionized bioinformatics by providing online DNA alignment of queries against large databases of reference genomes. However, assemblies constitute only 17% of archived bacterial data (110,898 assemblies versus 554,680 raw read datasets in the European Nucleotide Archive (ENA) as of October 2017) and generally only a fraction of those are indexed for BLAST search. Indeed, although high quality reference genomes are the gold standard, these are unachievable with short read data ^10-12^, and bad references discard or confound data which may be of interest. More fundamentally, many sequence datasets contain populations rather than clonal isolates, for which haploid assembly is a fundamentally poor summary for such data. As the archives continue to scale-up, it becomes critical to be able to rapidly filter them down to small datasets for subsequent analysis. The core requirement is therefore to be able to search heterogeneous historical and modern data, whether assembled or not, for presence of arbitrary sequence. In this study, we combine computational techniques previously used in web-search, with knowledge of bacterial population genetics, to develop a data structure, the *Bitsliced Genomic Signature Index (BIGSI)*, that solves this problem.

The first scalable search engine for raw sequence was developed in 2016, the Sequence Bloom Tree (SBT)^13^, a k-mer (fixed-length DNA word) index, developed to enable detection of specific transcripts in RNA-seq data. However, SBT and its recent successors^14-16^ shared with previous similar methods (Cortex, vari, McCortex)^17-20^ a scaling dependence on the total number of k-mers in the union of datasets indexed. For species where there is considerable k-mer sharing between datasets (e.g. human), this works well. However, bacteria are fundamentally different: even within a species there can be enormous diversity due to horizontal transfer of DNA (Supplementary Figure 1), and we will show that this diversity renders previous methods unable to scale. We remove this limitation, and use BIGSI to index the entire bacterial and viral content of the ENA as of December 2016 (447833 datasets; 170 Tbytes of data), using four orders of magnitude less storage than previous methods. This is the first time the archives have been made accessible to search, and we make a version publicly available at http://bigsi.io. We demonstrate applications to basic biology and surveillance: ultra-fast search for the colistin resistance genes MCR1-3, mapping host ranges of 2827 plasmids, and plotting the changing prevalence of antibiotic resistance mutations and genes in the archives.

## High accuracy queries with a lossy compressed DNA index

We developed a data structure suitable for storing bacterial genomic data, the *bitsliced genomic signature index* (BIGSI). We use the generic term “dataset” to refer to either assembled genomes or unassembled sequence read-files from clonal or non-clonal samples. BIGSI combines a k-mer index with constraints on sequence queries, described below. A bloom filter is a tool in computer science^21^ which stores data (here, k-mers) in a bit-vector (array of zeroes and ones) and answers set-membership queries (“is this k-mer contained in the set?”) probabilistically. The false negative rate is zero, and the false positive rate is controlled by two parameters (size of bit-vector and the number of (hash) functions used to generate the binary encoding, see Figure 1), creating a trade-off between false-positive rate and compression. We describe how we set the bloom filter parameters below. The BIGSI encodes data as a matrix where each column is a bloom filter of the k-mers in a dataset. Figure 1 shows schematics of how data is processed and stored, and details are in Methods. Note that incorporating a new dataset simply requires a new column is added, without needing to rebuild the index. Our implementation supports both disk-based and in-memory stores; all measurements reported are from the disk-based store. Although we use this as a k-mer index, it can be viewed as a probabilistic coloured de Bruijn graph^17,22^.

**Figure 1:**
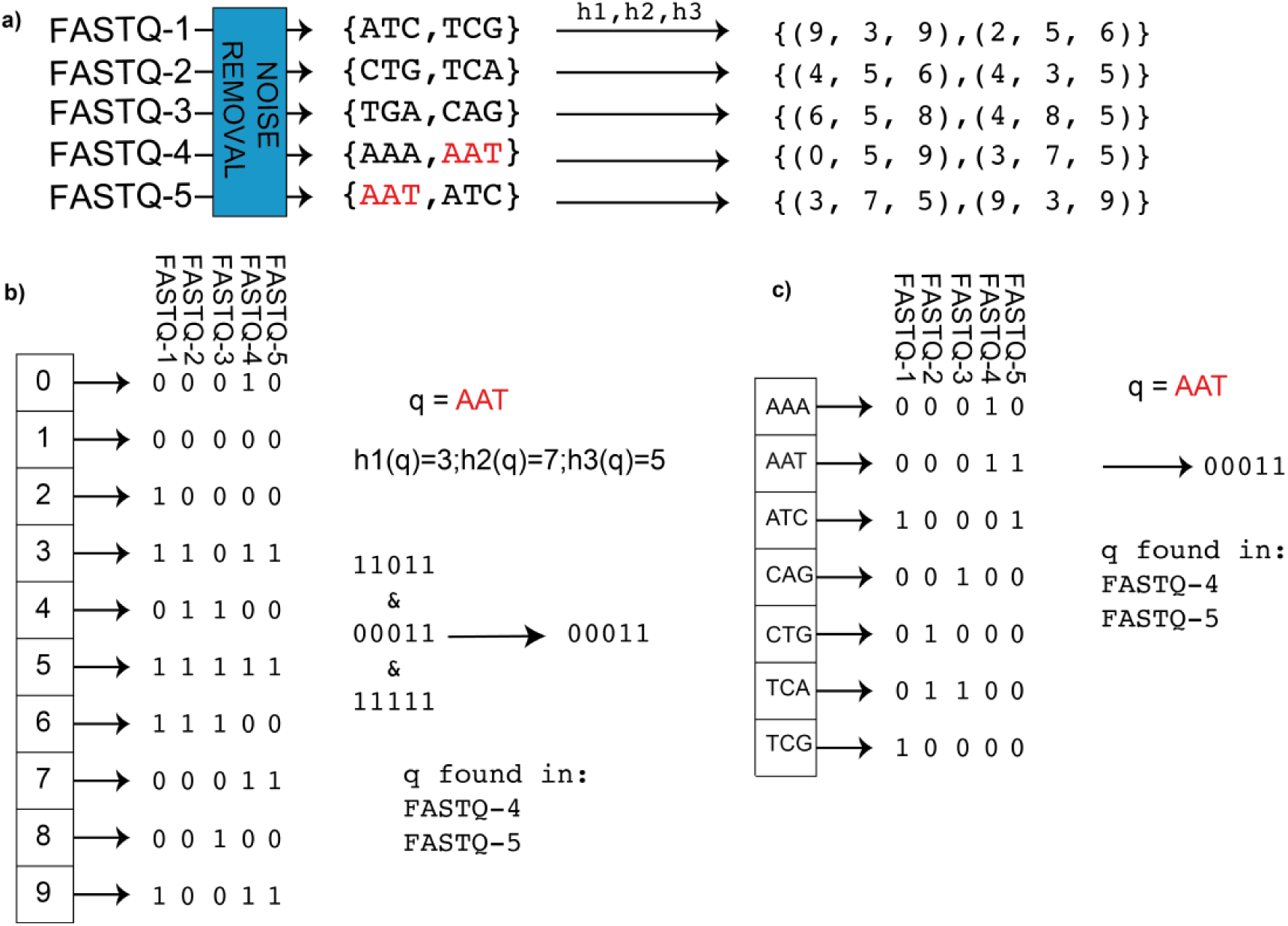
BIGSI encoding compared with naïve approach. a) BIGSI step 1: each input dataset (could be raw sequence data (FASTQ format) or assembly) is converted to a non-redundant list of kmers (with an optional denoising step to remove sequencing errors, detailed in Methods). A fixed set of *η* hash functions (*h*_1_, *h*_2_,…) is applied to each k-mer (*η* =3 in this figure), giving a tuple of positions which are all be set to 1 in a bit-vector (a Bloom Filter). b) BIGSI step 2. Each dataset is stored as a fixed length bloom filter, as a column in a rectangular matrix. To query the BIGSI for k-mer AAT, the *η* hash functions are applied to the query k-mer, returning *η* rows to be checked (namely 3,7,5 here). All columns (datasets) that have 1 in all of those *η* rows contain the query k-mer: these rows that are checked are called “bitslices”. Adding a new dataset requires just adding a new column. c) Naïve encoding for contrast. A complete list of all k-mers in all datasets form the rows of a large matrix, and columns are datasets. For any given k-mer, entries are set to one for datasets containing that k-mer. When a new dataset is added, the matrix grows vertically (new k-mers added) and horizontally (new column for new dataset).

To genotype a sequence, we query the index for all the k-mers within it. Exact matching requires all k-mers be present (threshold T=100%), and can be implemented as a fast AND operation on bit-vectors. Inexact matching, primarily used for long alleles, requires the presence of some proportion (T<100%) of k-mers be present, and is slower (See Figure 3a). Although BIGSI does not do an alignment, an approximation to a mega-BLAST alignment score can be inferred from the presence/absence pattern of kmers in the query; e.g if the k-mer size is 31, each SNP difference causes a window of 31 absent k-mers (details in Methods). We show in Supplementary Figure 2 the strong correlation (r=0.998) between Mega-BLAST score and BIGSI score for 100 *E. coli* AMR genes using a BIGSI of RefSeq-bacteria (release 81).

**Figure 2:**
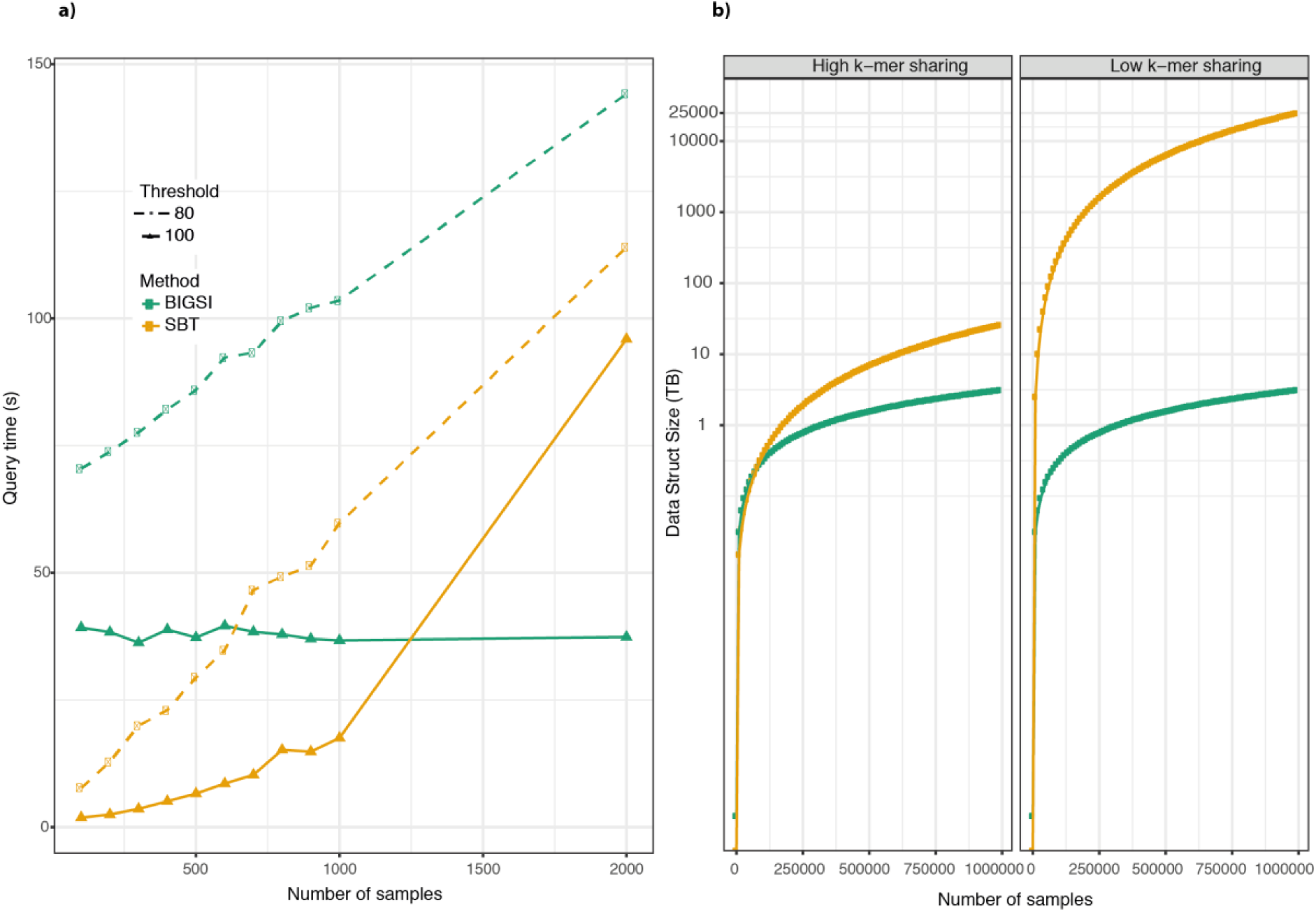
Scaling properties of BIGSI. a) Query times for 705 antimicrobial resistance genes when searching in a SBT (yellow) or BIGSI (green) for exact (T=100%, full lines) or inexact (T=80%, dotted lines) matches. Query times for BIGSI exact searches (T=100%) do not noticeably increase on this scale, as a result of the bit vector optimization possible for exact matching. b) Simulated scaling to 1 million datasets of peak data-structure storage requirements of BIGSI and SBT, comparing performance with high/low proportion of sharing of k-mers between datasets (note y axis is on log scale). In the high k-mer sharing regime only 100 new k-mers are introduced per dataset, whereas the low k-mer sharing regime introduces 10,000 new k-mers per dataset. Since BIGSI scales linearly with number of datasets and independently of the number of k-mers, it uses the same storage per dataset in each regime. However, SBT scales super-linearly with N since its bloom filter size depends on the total number of k-mers. For 1 million genomes with low kmer sharing (right), which is the case we care about for global indexing, BIGSI would use 3.1 Terabytes whereas SBT would use 25 Petabytes. When we index the ENA below, we find each dataset adds 100,000 new k-mers on average, 10x more than the low kmer-sharing regime simulated here, which would further penalize SBT.

**Figure 3:**
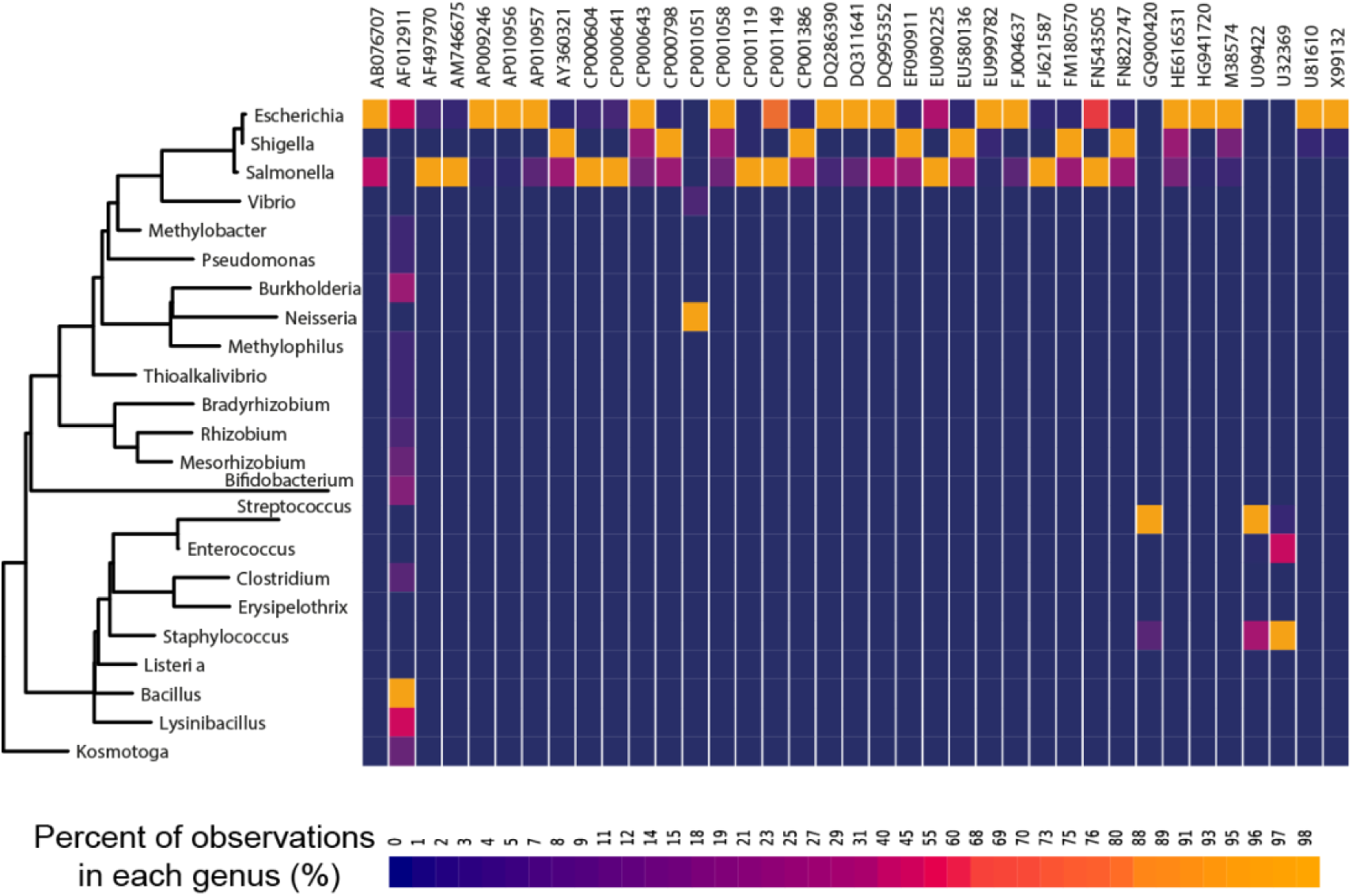
37 plasmid sequences found at least 5 times in more than one genus in the all-microbial-index. The heatmap shows the frequency of each plasmid within each genus; the tree on the left is a maximum likelihood rRNA tree of bacteria. The plasmid at the left (AF012911) with extremely wide phylogenetic distribution is a known cloning vector. The large amount of sharing between *Escherichia, Salmonella* and *Shigella* is consistent with known promiscuity within *Enterobacteriaceae*.

To search for a single nucleotide polymorphism (SNP), we create a sequence for each allele, with k-mer -1 bases on either side. By requiring multiple k-mers in the sequence be present, we reduce the false positive rate for SNP allele detection exponentially. Indexing at a smaller k-mer (31) than our minimum query length (61) enables both compression and a low error rate. The theoretical false discovery rate for an SNP allele from a probe (flanks plus allele) of length (2k-1) with bloom filter parameters is 10^-15^ per column (see methods) - well below the expected error rate from the underlying sequence data.

We measured query speed by first building a BIGSI of 3,480 datasets of *Mycobacterium tuberculosis* obtained from^23^, and genotyping 68,269 SNPs. Searching all datasets for these SNPs took just under 90 minutes on a single CPU core - an effective genotyping rate of above 46,000 genotypes per second.

We validated SNP genotyping accuracy using a subset of 100 of the *M. tuberculosis* datasets for which we had high quality SNP calls using samtools^24^ (see Methods). The concordance between methods was 99.997% with a total of only 286/682,690 discrepancies. We measured accuracy of longer allele detection by searching (with T=70% match) for a catalogue of *E. coli* Multi Locus Sequence Type (MLST) alleles and choosing the best scored allele for each gene. We then compared calls on a set of 954 datasets with the MLST allele calls from a high-quality caller: SRST2^25^. Where both methods made a call (6483/6678 alleles), there was 99.9% agreement; otherwise SRST2 failed (n=167), or BIGSI failed to find an allele version above T=70% (n=28).

## Scaling to millions of bacteria

We benchmarked the empirical scaling properties of BIGSI against the Sequence Bloom Tree (SBT)^13^ on a dataset of 2000 bacteria of the taxon *Enterobacteriaceae* from the ENA, comparing build and query times, and peak storage requirements on 12 increasing subsets ranging from 100 to the complete 2000 samples. For the full dataset, SBT required 4.9 TB of intermediate storage to construct the index and took 3 days to build. In contrast, BIGSI required 6.2GB space and took 5 hours to build – 784x smaller and a 14x faster build time. For each subset, we queried the SBT/BIGSI for 705 antimicrobial resistance genes with an average length of 847 bp and total query length 597,753bp. Exact and inexact (T=80%) query times can be seen in Figure 2a. SBT and BIGSI returned identical hits from the exact match search. In the inexact search, there was only one discordant query, where BIGSI returned two extra hits, both close to the threshold of 80%, that were not returned by SBT. This was likely due to the different construction of the underlying bloom filters as they are both found by SBT when the threshold was lowered to 70%. Both methods inexact query time scaled similarly with number of datasets. Constructing SBTs with larger numbers of datasets quickly becomes prohibitive in both storage and construction time required. We did not compare with the successors to SBT as they either have even higher intermediate disk use than SBT^14,15^, or were published as this paper was being finalized^16^ (November 2017; see Methods).

Finally, we simulated the scaling of storage requirements required to construct a SBT, and BIGSI for data sizes up to 1 million genomes in two regimes: firstly, for genomes with very high proportions of kmer-sharing, and secondly to species with lower proportions of kmer-sharing (e.g., most bacteria) - see Methods for details, and Figure 2b). BIGSI scales linearly with number of datasets, performing identically in both cases. In the low-kmer sharing regime (which is our focus) SBT would require 4 orders of magnitude more storage than BIGSI to construct (25 Pb rather than 3.1 Tb).

## Indexing all bacterial and viral WGS data

We set out to construct a BIGSI from all bacterial and viral WGS data-sets in the ENA “pathogens endpoint” (which contains all bacteria, all viruses and some eukaryote parasites, totaling 469,654 datasets). After excluding the eukaryotic genomes on the basis of size (see Methods), we were left with 447,833 datasets. The entire index required 1.5TB of storage, <1% of the original data size (170 TBytes) and contained more than 60 billion unique k-mers. Data download took 6 weeks, constructing bloom filters on the fly. Combining the bloom filters afterwards took approximately 2 days. We estimate that the intermediate storage required to build an SBT of the same data would have been >6.7PB.

We base a number of large-scale analyses below on this index, which we refer to as the all-microbial-index. In order to make statements about which genus mobile elements or alleles are found in we estimated the species and abundances present in each dataset of the all-microbial-index using the Bayesian abundance estimator Bracken^26^ (see Methods). We found over 90% of the datasets were from just 20 genera, and 65% were from the top 5 most common bacterial genera (*Salmonella, Streptococcus, Staphylococcus, Escherichia* and *Mycobacterium*); counts for the most prevalent bacterial genera are shown in Supplementary Figure 3.

## Application 1: ultrafast gene search

As a practical example, we searched for exact matches of the colistin resistance genes MCR-1/2/3, subject of intense scrutiny over the last 2 years^27-31^ across all 447,833 datasets in the all-microbial-index. Searching for all 3 genes took 1.73 s seconds in total, scanning 10x more genomes than previous publications. MCR-2 was not present, but we found MCR-1 in 169 datasets of 3 species *(E. coli, S. enterica, E. aerogenes)* and MCR-3 in 34 datasets *(E. coli, S. enterica, K. pneumoniae)* (see Supplementary Table 1).

## Application 2: estimating the host-range of plasmids and conjugative systems

We took 2827 plasmids from the ENA (see Methods, and Supplementary Table 2) and ran an inexact (T=40%) search for these in the all-microbial-index. We filtered these for hits with T>90% for downstream analysis. The total length of query sequence was 227 Mbp, and the query took 2120 CPU hours (11 days real time) on a single server using 8 cores and 1.5 Gb RAM per process. The search returned 665,619 hits with 121,758 unique accessions across 258 genera. Since contamination could confound observations of a plasmid in a genus, we excluded from this analysis (Application 2) all datasets containing evidence of more than one genus at abundance above 0.1%. Only 41% (=184652) of datasets and 62% of search hits passed this filter.

We often identified plasmids shared by closely related genera, notably among *Escherichia, Shigella* and *Salmonella,* and among *Enterococcus* and *Streptococcus.* We found 37 plasmids present in at least five datasets of at least two genera (Figure 3, Supplementary Table 2 & 6); and 5 in multiple orders and families.The plasmid pETHIS-1 (entry: AF012911) was found in 5 phyla, 10 taxonomic classes, and 17 genera. This plasmid is used as an expression vector and its identification in so many species serves as a positive control, confirming that BIGSI can spot similar plasmids across the database. Of more biological interest, the Tn916 conjugative transposon encoding tetracycline resistance that was first found in *Enterococcus faecium* and known to have broad host range^32^ (entry: U09422) was found in Streptococcus (n=3951), Staphylococcus (n=1212), Enterococcus (n=43), Clostridioides (n=29), Listeria (n=19) and Erysipelothrix (n=11).

Sampling biases in the ENA preclude inference about plasmid prevalence, but they do allow us to test if plasmids bearing antibiotic resistance (ABR) genes are more widely phylogenetically distributed than those bearing none. We used a large subunit ribosomal RNA tree to compare genera. We defined “phylogenetic spread” of a plasmid as the median of the patristic distances between all pairs of genera in which the plasmid is seen. We test whether plasmids with at least 3 ABR genes (abbrev. 3-ABR) are more widely distributed across the phylogeny than those with none, by comparing the two phylogenetic spread distributions (Figure 4). We find the 3-ABR plasmids are indeed more widely spread (p=0.0024, permutation test, see Methods and Supplementary Figure 4). Given the underlying data, we are most confident of this conclusion within the *Enterobacteriaceae*, beyond which we would want to replicate this with wider sampling of the phylogeny.

**Figure 4:**
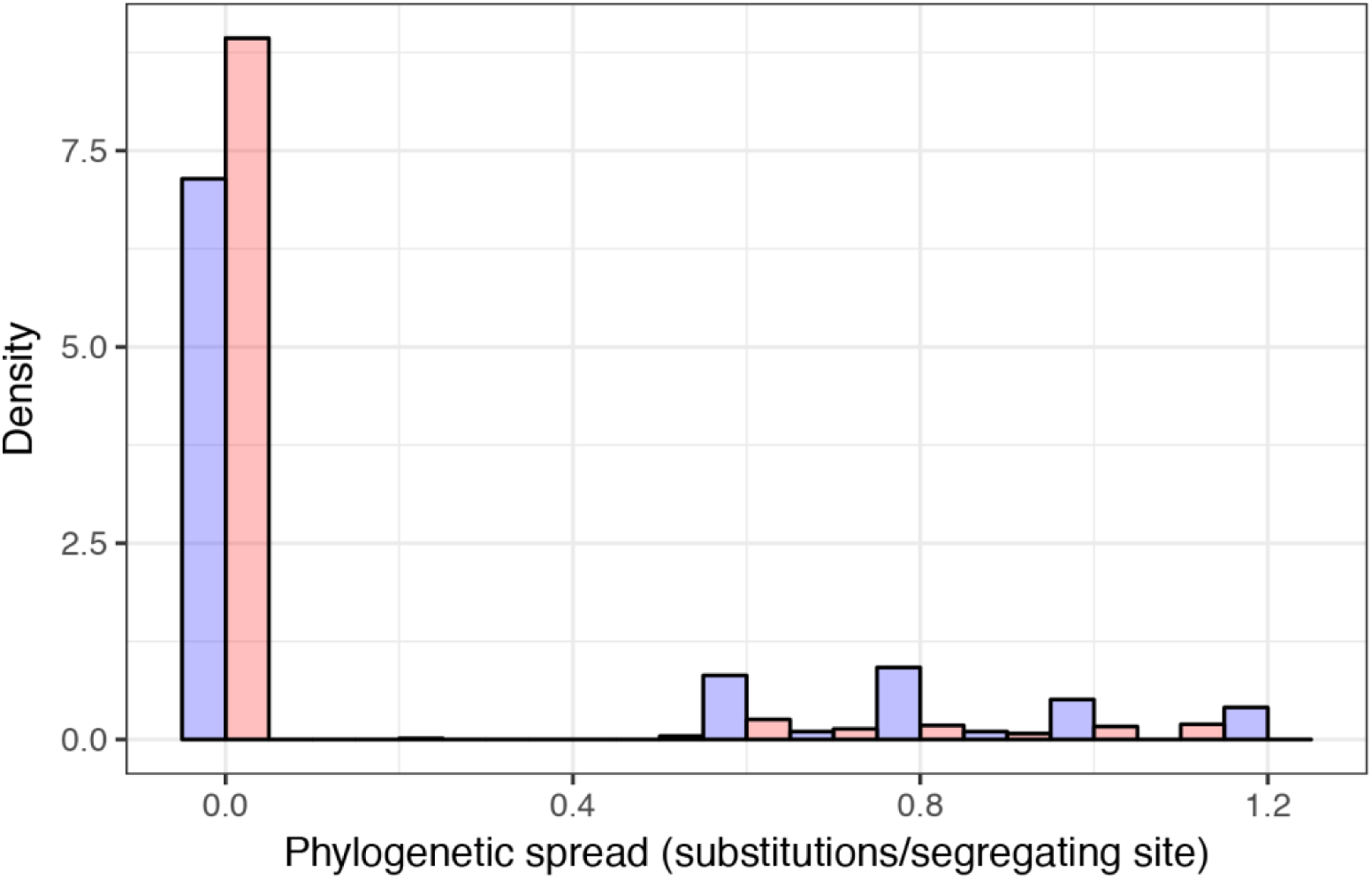
Comparison of phylogenetic spread (median of pairwise distances between all pairs of genera in which a plasmid is seen) of plasmids containing at least 3 antibiotic resistance genes (n=98, purple) with those bearing none (n=665, peach) – histograms are normalized to allow comparison (probability densities). Distance measured on the large subunit rRNA tree from SILVA. Units of phylogenetic spread are substitutions per site.

The distribution of conjugative systems for transfer of DNA between bacteria was previously analysed in 1124 genomes^33^ using sensitive, but slow, protein profiles searches. We extended this analysis to the whole ENA, searching for exact matches in the all-microbial-index for the previously identified key components of these systems (relaxases (MOB) and type 4 secretion systems). Of the 184,652 datasets, 36030 (19.5%) had a putative conjugative system-consistent with the previous estimate of 18%. This proportion varied by phylum, from 0.5% in *Spirochaetes* to 31.7% in *Firmicutes* (see Table 1, Supplementary Figure 4). At a finer scale these observations provide valuable information on the potential spread of antibiotic resistance genes. For example, focusing on datasets with relaxases of the type MOBT we observe genetic flux between *Staphylococcus* and *Streptococcus*, but not with *Salmonella,* and this information facilitates the assessment of the risk of spreading between taxa. This flux does not need to strictly follow phylogenetic lines. For example, we observed MOBQ in *Salmonella* and *Streptococcus* but not *Staphylococcus*, indicating a different probability of cross-genus (and cross-phylum) transfer by conjugation (data in Supplementary Table 4).

**Table 1:**
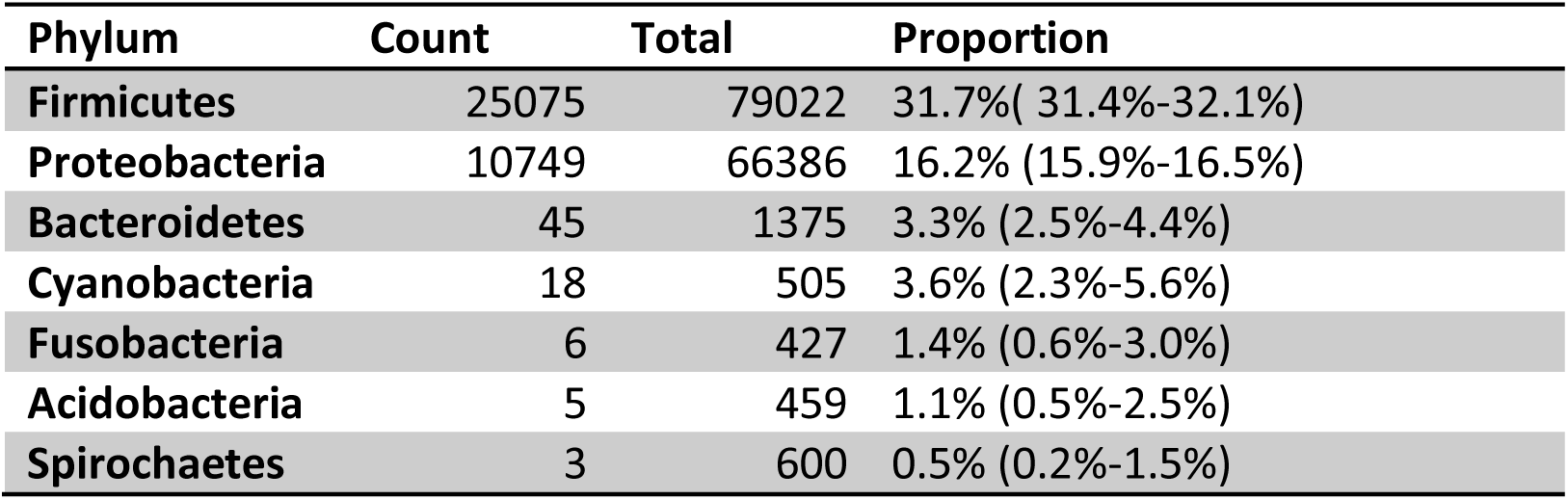
Observations of putative conjugative elements across phyla. The counts of phyla with search hits for putative conjugative elements and the counts of all samples from each phylum in all-microbial-index. Dividing the ICE hits in the all-microbial-index by the counts of members of a phylum in all-bacteria is an estimate of the proportion of samples of each phylum-containing conjugative system. Confidence intervals are calculated using the Wilson interval.

## Application 3: growth of antibiotic resistance prevalence in the archives

Monitoring of antibiotic resistance burden is a significant goal that is currently unachievable, in particular because we discover new resistance genes and then need to search the entire back-catalog of genomes (such as with MCR1-3). We sought to prove the concept by running a typical scan across the ENA. We downloaded all 2157 sequences associated with antibiotic resistance from the CARD database (v1.1.7)^34^ and searched for these in the all-microbial-index with thresholds of 100% and 70%. The exact searches for a gene took on average 1.1s and returned 438 hits per queries gene, resulting in 944,862 hits in 193,582 unique accessions across 250 genera (full results in Supplementary Table 5). An inexact search (T=70%) on average took 34.4s and returned 5320 hits. We show in Figure 5a the increasing count of ABR genes in the archives, as a function of year of upload to the ENA/SRA. Restricting the analysis to *Staphylococcus,* we find (Figure 5b) the proportion of datasets containing *mecA* gene (causing methicillin resistance) dropping from a high of 70% in 2013 to 40% in 2016, during which period all the *tet* and *aac* genes also drop in prevalence. By contrast, essentially all resistance genes have gone up in prevalence in *Klebsiella* (Figure 5c).

**Figure 5:**
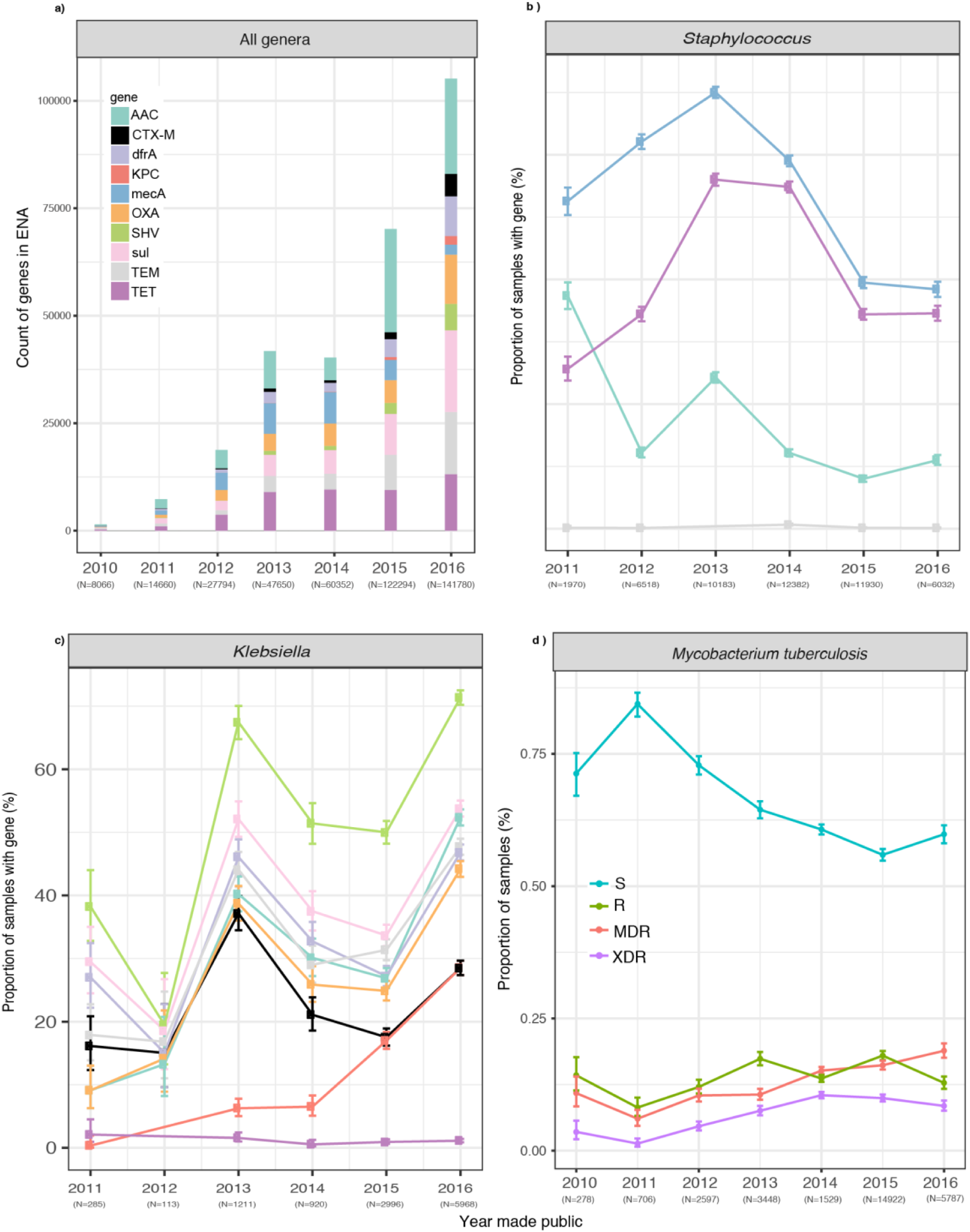
a) counts of samples in the all-microbial index containing a range of ABR genes; each gene treated independently, so a single dataset containing both CTX-M and OXA for example, will be counted twice b) Year-by-year frequency (defined by date of public availability) in *Staphylococci* (dominated by *S. aureus)* of *mecA,* and all *tet* and *aac* genes, which encode resistance to methicillin, tetracycline and aminoglycosides respectively. Archive-prevalence dropping for all since 2013. c) Year-by-year frequency in Klebsiella of various ABR genes; increase in prevalence since 2014 may be due to increased Extended Spectrum Beta-Lactamase surveillance and sampling of KPC resistant *Klebsiella* globally. d) Year-by-year breakdown of *M. tuberculosis* datasets, classified by genotypes as resistant (R), pan-susceptible (S), multiple drug resistant (MDR), extensively drug resistant (XDR) as follows. All datasets were genotyped for variants from the resistance catalog from^23^, then classified as resistant or susceptible to Datasets were classed as MDR (multi-drug resistant) if resistant to isoniazid and rifampicin, as XDR (extensively drug-resistant) if MDR and also resistant to a fluoroquinolone, and any of capreomycin, kanamycin and amikacin, and as Resistant if resistant to any antibiotic but not MDR or XDR, and susceptible otherwise.

In *M. tuberculosis,* resistance is driven primarily by amino acid mutations^1,35^. Genotyping all datasets in the all-microbial-index simultaneously (of which 30,226 were *M. tuberculosis)* for the 206 resistance mutations from Walker et al. ^23^ took 103 minutes on a single-core, around 10,000x faster than typing each dataset individually with the fast resistance prediction software, *Mykrobe predictor^1^.* The results show (Fig 5d) a rise in prevalence in the archive of MDR-TB since 2011 to a peak of 18.9% in 2016.

## Discussion

There is an urgent need for a global infrastructure for surveillance and management of infectious disease - microbes know no borders, and evolve faster than we modify our responses^36,37^. Vital analytic and visualization tools for SNP-based analyses are being developed in response to emerging viral outbreaks^38^, but the problems of scalability, more variable genomes and mobile elements have not been addressed. We foresee a world where millions of bacterial and viral samples have been sequenced and shared, some from very controlled and high quality clinical and public-health sources providing high-value metadata (e.g. the open Genome Trakr database of food-borne pathogens in the USA), and others of varying provenance. A scalable online sequence search facility would be critical to this endeavor. It would provide data not just for urgent outbreaks, but also for monitoring global strain, plasmid and resistance prevalence in humans, animals and the environment. We have demonstrated the core operations needed for such a search tool, on a scale never before achieved, indexing the entire bacterial and viral WGS content of the global DNA archive. Since new datasets can be rapidly appended to the index, the ability to grow incrementally as new datasets are sequenced is guaranteed. Finally, the method is ready for a future where finished reference genomes become routine, as the index works equally for both raw data and assemblies.

Searching the DNA archive is one example of a “document retrieval” problem, a subject which has been intensely studied and successfully implemented at massive scale by internet search engines. Our “search terms” are k-mers from SNPs/alleles, and our “web-pages” are raw read datasets or assemblies (see Figure 6). Similar approaches to ours (also using bitsliced signatures) have been used for text search^39,40^, but were largely abandoned after Zobel et al showed in 1998 that an alternative method (inverted indexes) performed better for natural language^41^. One notable exception this year was has been Microsoft Bing search engine^42^, which also revives them. For our use case, where each new bacterial dataset brings new variation, bitsliced signatures provide much better scaling than inverted indexes. Web and (microbial) DNA search have different dimensionality, as the language of bacterial genomes is vastly more complex than English. Our dataset was only 10^6^ documents but contained 10^10^ unique words, and this would continue to increase with more data, whereas Google indexes 10^12^ documents containing (we estimate) 10^8^ words with a much more slowly growing lexicon. As a result, we expect fruitful future interactions between the genomic and document retrieval communities.

Google revolutionized the utility of the internet, and now search is intrinsic to how we interact with the online world. BLAST did something similar for molecular biology in the 1990s, but does not scale even to the current archive. We are currently investigating implementing the BIGSI as a live service at the EMBL-EBI, updated as data is added to the ENA. We believe our approach, and improvements that will surely follow, will put our shared DNA store at everyone’s fingertips.

## Online Methods

### BIGSI construction and querying

BIGSI indexes a set of N (number of datasets) bloom filters by position in the bloom filter. Each bloom filter must be constructed with the same parameters (m, *η),* where m is the bloom filter’s length in bits and *η* is the number of hash functions applied to each k-mer. The same hash functions must also be used to construct each bloom filter. To construct a BIGSI, the N bloom filters are column-wise concatenate into a matrix. The row index and row bit-vectors are then inserted into a hash table or key-value store as key-value pairs so that row lookups can be done in O(1) time. This set of key-value pairs can be stored on disk, in memory or distributed across several machines. To insert a new bloom filter we simply append it as a column to the existing bitmatrix. To query the BIGSI for a k-mer we hash the k-mer *η* times, look up the resulting keys in the key-value store, and take the bit-wise AND of the resulting bit-vectors (See Figure 1).

### Parameter choices

The choice of BIGSI parameters (m, *η*), depends on: the maximum number of k-mers expected in any dataset (*K_max_*), the number of datasets (N) expected, the shortest length of the query sequence to be supported (*L_min_*), the k-mer size (k) and the maximum number of acceptable false discoveries per query (*q_max_*). Since each query of length (L) will consist of 𝓛 = L-k + 1 k-mers the expected number of false discoveries (V) for any query can be calculated as *q* = *E*[*V*] = *Np^𝓛^* where p is the false positive rate of the bloom filter. Parameters m and *η* determine false positive rate for a bloom filter with *K_max_* elements – if there are fewer elements, then the false positive rate will be lower. We assume below that all bloom filters have the maximum number of k-mers inserted to give an upper bound on error rate.

To keep q below a chosen threshold *q* < *q_max_* for a given N and k, p must be chosen to satisfy q for the shortest query of length *L_min_*:

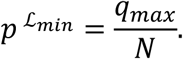

Therefore, the desired bloom filter false positive rate is

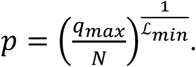

Since, for a given number of inserted kmers (n), and desired false positive rate (p), optimal bloom filter parameters can be determined by the following formulae^43^

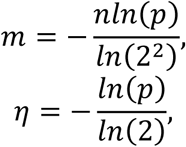

which becomes:

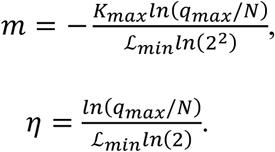

For example, given

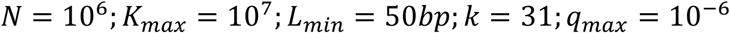

the resulting expected number of false positives per kmer-lookup per bloom filter (p), would be:

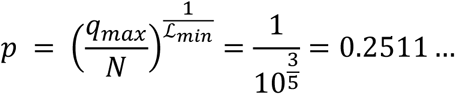

Solving the above equations gives:

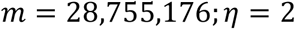

### BIGSI parameters for all-microbial-index

We assume initially that a bacterial dataset contains at most 10 million k-mers since bacterial genomes are generally under 6 Mb in length, leaving 4 million k-mers available for some sequencing errors which escape de-noising, and genomic variation. Unless otherwise specified we use BIGSI parameters m=25,000,000, *η* = 3 for all analyses. For these parameters, the upper bound on number of false discoveries, *q_max_* for an SNP allele from a probe (flanks plus allele) of length (*L_min_* =2k-1=61) is 10^-9^ (if *K_max_* = 10^7^,N = 10^6^).

### Code availability

An open source (MIT license) implementation of BIGSI can be found at https://github.com/phelimb/BIGSI. BIGSI v0.1.2 supports both disk-based (via Berkeley-DB in memory (via python dictionaries), and distributed in memory (via redis (https://redis.io), as used in Twitter) key-value stores. All the results discussed below use the Berkeley-DB, disk-based backend.

### Public instance

We have made a public instance of our index of the ENA available at http://bigsi.io, where the user can paste sequence and search. The user receives a list of accessions containing the given query (which can be clicked on to get to the ENA webpage for the accession), and also the species abundance estimates from Bracken. This instance uses the redis in-RAM implementation and is hosted by CLIMB^44^ on a 3Tb RAM server.

### Scoring of BIGSI queries

BIGSI search hits can optional be scored using an approximation to the ungapped alignment scoring scheme used by megaBLAST. To do this, we take the presence/absence vector for a query of 𝓛 k-mers. From this, we estimate the approximate number of mismatches of the query from the hit by counting the number of zeroes in contiguous runs of length greater than 1, and dividing by the k-mer size. From these estimated mismatches and matches we calculate a score for an ungapped alignment, with p-values calculated using the same scheme as BLAST. By default the costs are -2 for a mismatch and +1 for matched position.

### Comparison of query time and storage requirements of BIGSI and SBT

We randomly chose 2,000 *Enterobacteriaceae* cleaned de Bruijn graphs from the all-microbial-index accessions and we then further randomly sub-sampled these into collections of 10, 100, 200, 300, 400, 500, 600, 700, 800, 900, 1000, 2000 datasets. Jellyfish v2.2.5 was used to count the unique k-mers in the set of cleaned graphs. A BIGSI of each set of datasets was built with parameters (m = 2.5 × 10^7^;*η* = 3). A SBT was built for each set of datasets with *η* = 1 and bloom filter size (m) equal to the count of the total number of k-mers in the collections of graphs. Construction and query time analyses were run on a Dell PowerEdge R820 with 32 cores and 1 Tb RAM. For reproducibility we give the precise commands: the search time comparison was run with ‘bt query‘and ‘cbg search‘, using k-mer thresholds 100% and 80%. BIGSI was run in server mode (‘hug cbg‘) for the search profiling in order to exclude python boot up time overhead.

Simulation of storage requirements for a BIGSI for N datasets is given by:

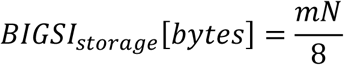

We model the storage for SBT as follows. Although it is possible to append to an SBT incrementally, as new microbial datasets will keep adding new k-mers, this will lead to saturation of the root-level bloom filter in the SBT, and a collapse in query performance. This can be avoided by reconstructing the SBT, ensuring the bloom filters are large enough to support the full set of k-mers. Thus, since the best case for a binary tree with N leaves is 2N-1 nodes, we estimate:

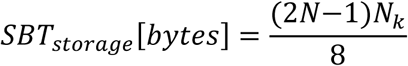

where *N_k_* is the total number of k-mers in the combined set of datasets and also equal to the size of the bloom filter required.

We simulated two regimes. For high-kmer sharing, we assumed each new dataset added 10,000 new k-mers, which corresponds

### No benchmarking of successors to SBT

The split sequence bloom tree (SSBT)^14^, the allsome bloom tree^15^ and Mantis^16^ are more recent improvements on the SBT, focussed also on human transcriptomes. SSBT and Allsome modify their bloom filter trees compared with SBT, to capitalise on shared sequence between datasets. Although the ENA content is heavily biased towards a few genera, the amount of sharing between datasets is still much lower for these bacteria (several phyla) than for humans (one species), so the compression benefit will be lower. Pragmatically, both SSBT and Allsome have larger intermediate uncompressed indexes than the SBT, and so we did not benchmark against them.

Mantis was published as we were finalising this paper (Nov. 2017), and so we have not benchmarked against it. As currently implemented, the data structure does not support incremental insertion, as it needs up-front the full set of k-mers and for each k-mer, the list of datasets containing it (“colour class”, stored as a bit-vector). However, this is not a fundamental limitation, and a (less efficient) iterative construction algorithm is known, though not implemented (pers. comm. Rob Patro). The scaling properties of the final index for Mantis (for bacterial data) are likely to be better than SBT, SSBT and Allsome, since it can use compression on the colour class bit-vectors, avoiding a quadratic scaling of the colour-class matrix. Scaling will be superlinear but the exponent is data dependent, and unknown. Finally, there is currently an intermediate stage where the uncompressed colour-class matrix is held in RAM, scaling quadratically in number of datasets, which again could be removed by reimplementation.

### Using an array of BIGSI for longer genomes

A limitation of BIGSI is that *K_max_*, the maximum number of k-mers per dataset, must be set in advance. One way to extend BIGSI to datasets with varying k-mer cardinality is to build a nested structure of multiple BIGSIs with different *K_max_*, e.g. *K_max_ =* 10^5^,10^6^,10^7^, etc …, and insert each sequence into the appropriate level by k-mer counting before insertion.

### Genotyping accuracy measurement on TB

Conservative SNP calls were made using Cortex^17^ (independent workflow, k=31) on 3480 *Mycobacterium tuberculosis* datasets from Walker et al^23^. Singleton variants were discarded, and a non-redundant list of 68,695 SNPs was constructed. We generated “probe sets” consisting of a reference and alternate alleles of these variants from the NC_000962.3 reference. An index of the 3,480 datasets was built and 100 random datasets were genotyped at the 68,695 sites as follows: Each allele of the probe-set is searched for in the BIGSI resulting in Boolean presence/absence of each allele. If only a reference allele is present the genotype is returned as 0/0, if only an alternate allele 1/1, if both 0/1 and if neither -/-. We compared the concordance of genotypes of the 100 random datasets with those generated with the samtools^24^ pipeline from Walker et al^23^, excluding filtered positions. As described in the main text, the concordance between methods was 99.997% with a total of 286/682,690 discrepancies. The majority of these discrepancies (203/286) were mixed (heterozygous) calls from BIGSI; samtools had been run with a haploid model it did not make any mixed calls, so we expect some of these were correct.

### Indexing of ENA snapshot

The fastqs from accessions listed in Supplementary Table 6 were downloaded via ENA’s Globus FTP and included all WGS bacteria and viruses, but also eukaryotic parasites with larger genomes, which we did not intend to index. Most eukaryotic genomes were removed implicitly, since we set thresholds to exclude datasets with too many kmers for a 5Mb genome. De Bruijn graphs (k=31) were constructed and cleaned from the downloaded fastq files using mccortex^19^ v0.0.3-539-g22e27b7.

mccortex31 build -t 1 -m 7G -k 31 -s “DATASET_ID” -1 “FASTQ_FILES”

mccortex31 clean -m 7GB -B 2 -U -T

De Bruijn graph error cleaning and tip trimming were performed using mccortex. Bloom filters were built using the k-mers from the cleaned graphs with parameters (m = 2.5 × 10^7^;*η* = 3, *K_max_* = 10^7^) with BIGSI v0.2 as follows:

cbg init -k 31 -m 25000000 -h 3

cbg bloom -c “CLEANED_GRAPH_FILE”

Of the full set of datasets, 4.6% (21,822/ 469,654) fastq files failed to produce a resulting bloom filter. Of these 21,822: 7,799 exceed the maximum number of k-mers allowed after error cleaning (namely 10^7^) and 14,023 exceeded the maximum number of k-mers allowed in the raw dataset (namely ~7×10^9^).

The unique k-mers in the union of the cleaned graphs were counted using the redis (v3.2.6) hyperloglog (https://redis.io/commands#hyperloglog) approximate cardinality counter. In the union of all cleaned graphs there were 6.05 × 10^10^ ± 5 × 10^7^ k-mers. We estimate this number would have been at least an order of magnitude higher without the denoising step where mccortex removed sequencing errors.

### Species identification

The proportion of species in each dataset was determined using Kraken^45^ and Bracken^26^. Kraken v0.10.5 was run on the k-mers from each cleaned de Bruijn graph using the minikraken 20141208 database. The resulting taxonomy labels assigned by Kraken were then analysed by Bracken (commit version vfd88a06a) to estimate the proportion of k-mers originating from each species present in a dataset. Bracken failed to report species abundance for 12,889 datasets. The taxonomic data for the remaining 434,944 datasets is reported in Supplementary Table 6.

### Plasmid search and exclusion of contaminated datasets

2,826 plasmid sequences were taken from the ENA plasmid pages (www.ebi.ac.uk/genomes/plasmid.html; December 2016) (See Supplementary Table 2) and downloaded from the ENA. We then queried the all-microbial-index for these sequences with a proportion of k-mers threshold of 40% (T=40%) present and filtered for hits with T >= 90% for downstream analysis. Queries were run with 1 GB cachesize (memory) per process and parallelised across 8 vCPUs.

In order to determine distribution of plasmids across taxa, while avoiding ENA/SRA metadata errors, we filtered these hits for datasets which (were bacteria and) had no secondary genus above 0.1% frequency. This criterion was chosen to avoid multi copy plasmids from contaminating species establishing false positive hits within a non-host genus. 41% (184652/447,833) of accessions and 62% (415,181/668,720) of search hits passed this filter. We then filtered for all plasmids which had been seen at least 5 times each in more than one genus and had less than 99% of their observations in the most frequent genus. We found 37 plasmids across 13 genera matched these criteria. By simulating mixtures of *S. enterica* and *E. coli* at relative abundances of 0.0001, 0.001, 0.01, 0.02, 0.05, 0.1, 0.15, 0.2, 0.25, 0.3 we found we could observe the minority species above 2% frequency (the limit of detection was not lower because we had applied Kraken *after* error-cleaning of de Bruijn graphs). All 37 plasmids reported had at least one observation at a copy number of 5 (which, since we could detect contaminants at 2% frequency, would correspond to a copy number of above 250 if it came from a contaminant) and 16/37 had an observation at 2000x copy number.

### Phylogenetic spread of plasmids

We excluded contaminated samples as above, and plasmids with no hits in the all-microbial-index. We used the APE R package to calculate a patristic distance matrix between all genera in the Silva^46^ 23S rRNA-based phylogenetic tree built using RAxML with GTR-GAMMA distances (release s123_LSU, https://www.arb-silva.de/projects/living-tree/). For all plasmids, we took the N genera in which they were found and calculated the N-choose-2 distances between these using the above matrix. We then compared the cumulative distribution functions of the phylogenetic spread (since these intrinsically show how zero-centred the distribution is) using a permutation test, with test statistic the difference in 95%-quantiles. The 95% quantiles were 1.11 and 1.99 for non-ABR and 3-ABR plasmids respectively, and a permutation test with 1 million replicate permutations gave p=0.0024 – see Supplementary Figure 4 for the histogram.

### Identification of conjugative systems

The relaxases (MOB) of the previously described types (MOB_B, MOB_C, MOB_CQ, MOB_F, MOB_H, MOB_P,MOB_T, MOB_V) and the ket T4SS components (VirB4_TRaU, VirD4_TcpA) were used as defined in Guglielmini et al ^33^ and Supplementary Data 1 in the all-microbial-index with T=100%. Full search results are available in Supplementary Table 7. Results were filtered for bacteria and contamination following the same method as described in “Plasmid search”. Accessions with at least one MOB and the two key components ofT4SS were said to contain a putative conjugative system. BIGSI does not return copy number, or location on chromosome or plasmid, so it was not possible to determine if the genes were co-located on a chromosome or on a plasmid.

### MCR-1,2,3

We searched for MCR-1, MCR-2, MCR-3 in the all-microbial-index using kmer percent threshold T=100%. See Supplementary Table 1 for sequences and results.

### Searching for ABR genes in the ENA

We downloaded all 2157 sequences associated with antimicrobial resistance from the CARD database (v1.1.7)^34^. We searched for these in the all-microbial-index with thresholds of 100% and 70%, using a 1 Gb cache size, and 8 CPUs. A full table of the search results can be found in Supplementary Table 5.

### Searching for *M. tuberculosis* variants in the ENA

We searched the all-microbial-index for the variants from the catalogue described in ^23^ by generating “probe sets” consisting of a reference and alternate alleles of these variants from the NC_000962.3 reference and searching for these alleles. If only a reference allele is present the genotype is returned as 0/0, if only an alternate allele 1/1, if both 0/1 and if neither -/-. From the resulting genotypes, we classified each of the datasets as resistant or susceptible to 12 antibiotics following the model described in^23^. The date when this data was first available to the public was extracted from its ENA metadata. Datasets were classed as MDR (multi-drug resistant) if resistant to isoniazid and rifampicin, as XDR (extensively drug-resistant) if MDR and also resistant to a fluoroquinolone, and any of capreomycin, kanamycin and amikacin, and as Resistant if resistant to any antibiotic but not MDR or XDR, and susceptible otherwise.

## Data availability

All supplementary tables can be found at Figshare here: https://doi.org/10.6084/m9.figshare.5702776

## Author contributions

ZI, GM designed and oversaw the study, PB invented the method, developed software and performed analyses, HdB performed analyses for the plasmid study, EPCR co-designed and analysed the conjugative system and plasmid analysis, ZI wrote the paper, all authors made detailed feedback on the paper.

## Acknowledgements

We would like to thank, for critical reading and helpful suggestions: Martin Hunt, Jerome Kelleher, Janet Thornton, Rob Patro, John Marioni, Andrew Page, Simon Gog, Timo Bellman, Florian Gauger. For enormous assistance with data download from EBI: Robert Esnouf, Guy Cochrane. For hosting our BIGSI demonstration: CLIMB. We acknowledge funding from Wellcome Trust Core Award Grant Number 203141/Z/16/Z. ZI was funded by a Wellcome Trust/Royal Society Sir Henry Dale Fellowship, grant number 102541/A/13/Z. PB was funded by Wellcome Trust Studentship H5RZCO00. EPCR was funded by an ANR grant (MAGISBAC). GM was funded by Wellcome Trust grant 100956/Z/13/Z.

## Supplementary Figures

**Supplementary Figure 1.**
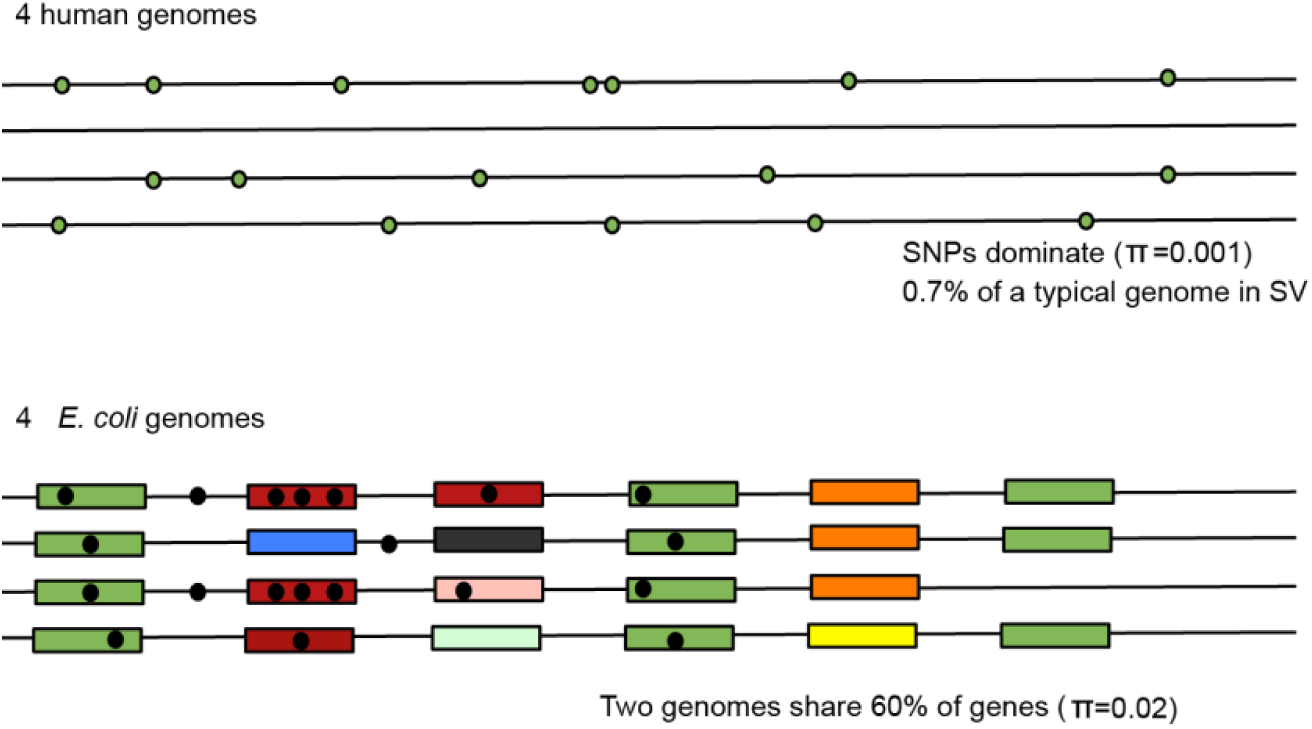
Cartoon comparison between human and E. coli genomes: Cartoon comparison of human genomes (above) and E. coli (below) as a representative bacterium. In humans, genetic variation is dominated by relatively sparse single nucleotide polymorphisms (SNPs), nucleotide diversity π=0.001, and less than 1% of a typical genome lies in a structural variant (SV) [1]. Human genomes are therefore relatively compressible. In stark contrast, genes make up around 88% of an E. coli genome [2], yet two E. coli genomes may only share around 60% of their genes [3], and conserved genes have much higher nucleotide diversity (0.02) [4]. Thus, bacterial genomes present different compression and indexing challenges to human genomes. 1. Genomes Project, C. et al. A global reference for human genetic variation. Nature 526, 68-74, doi:10.1038/nature15393 (2015). 2. Blattner, F. R. et al. The complete genome sequence of Escherichia coli K-12. Science 277, 1453-1462 (1997). 3. Touchon, Marie, et al. “Organised genome dynamics in the Escherichia coli species results in highly diverse adaptive paths.” PLoS genetics 5.1 (2009): e1000344. 4. Kaas, Rolf S., et al. “Estimating variation within the genes and inferring the phylogeny of 186 sequenced diverse Escherichia coli genomes.” BMC genomics 13.1 (2012).

**Supplementary Figure 2.**
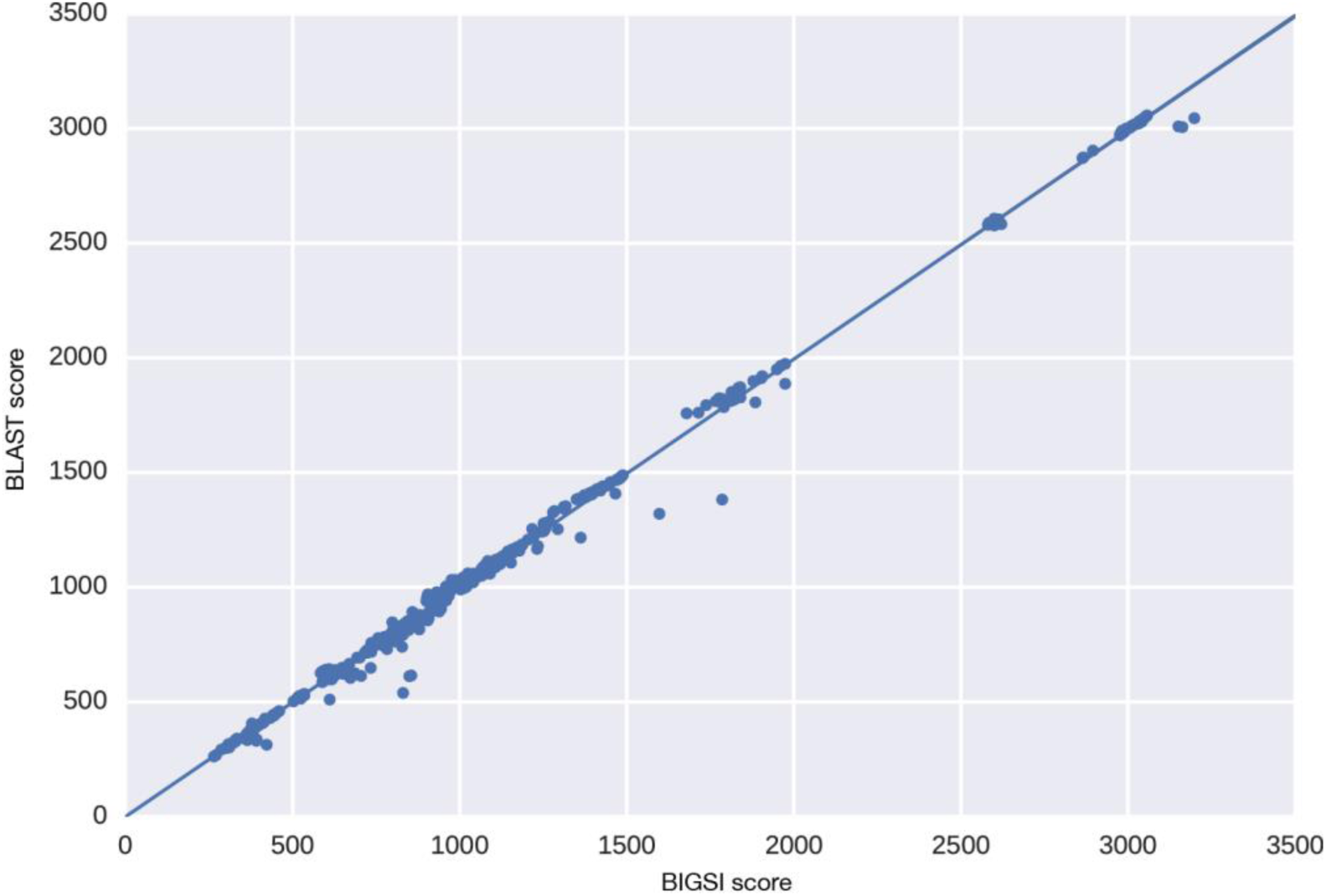
BIGSI scores vs megaBLAST scores:
megaBLAST scores for a search of 100 antimicrobial resistance genes in a BLAST database of RefSeq-81 vs the equivalent BIGSI scores in a search of a BIGSI of RefSeq-81. Pearson correlation of the scores was r-0.998.

**Supplementary Figure 3.**
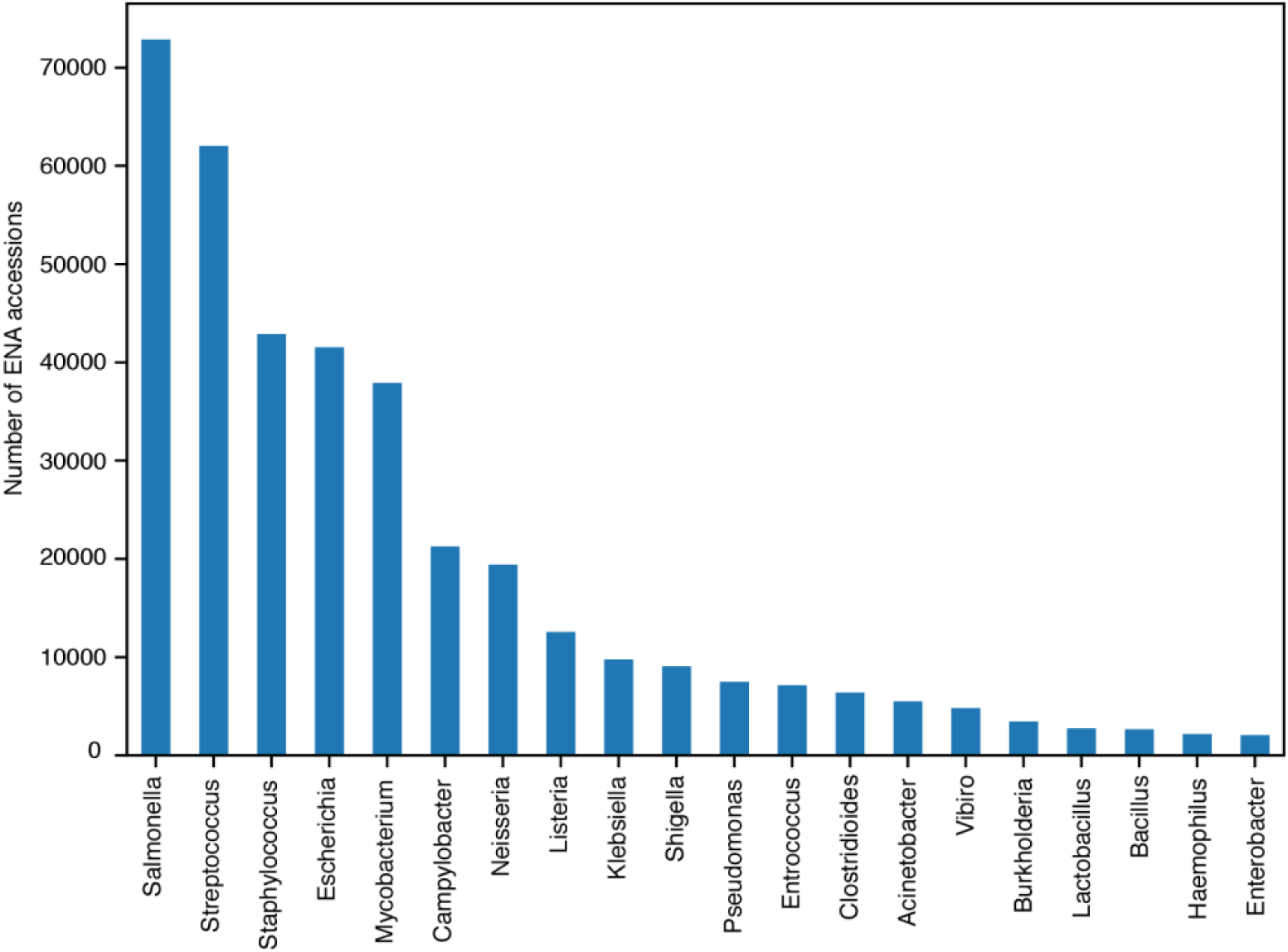
Counts of the most frequent bacterial genera in the SRA/ENA:
Counts of the most frequent bacterial genera in the all-microbial-index. Over 90% of the datasets were isolates of these 20 genera and 65% from the top 5 most prevalent genera.

**Supplementary Figure 4.**
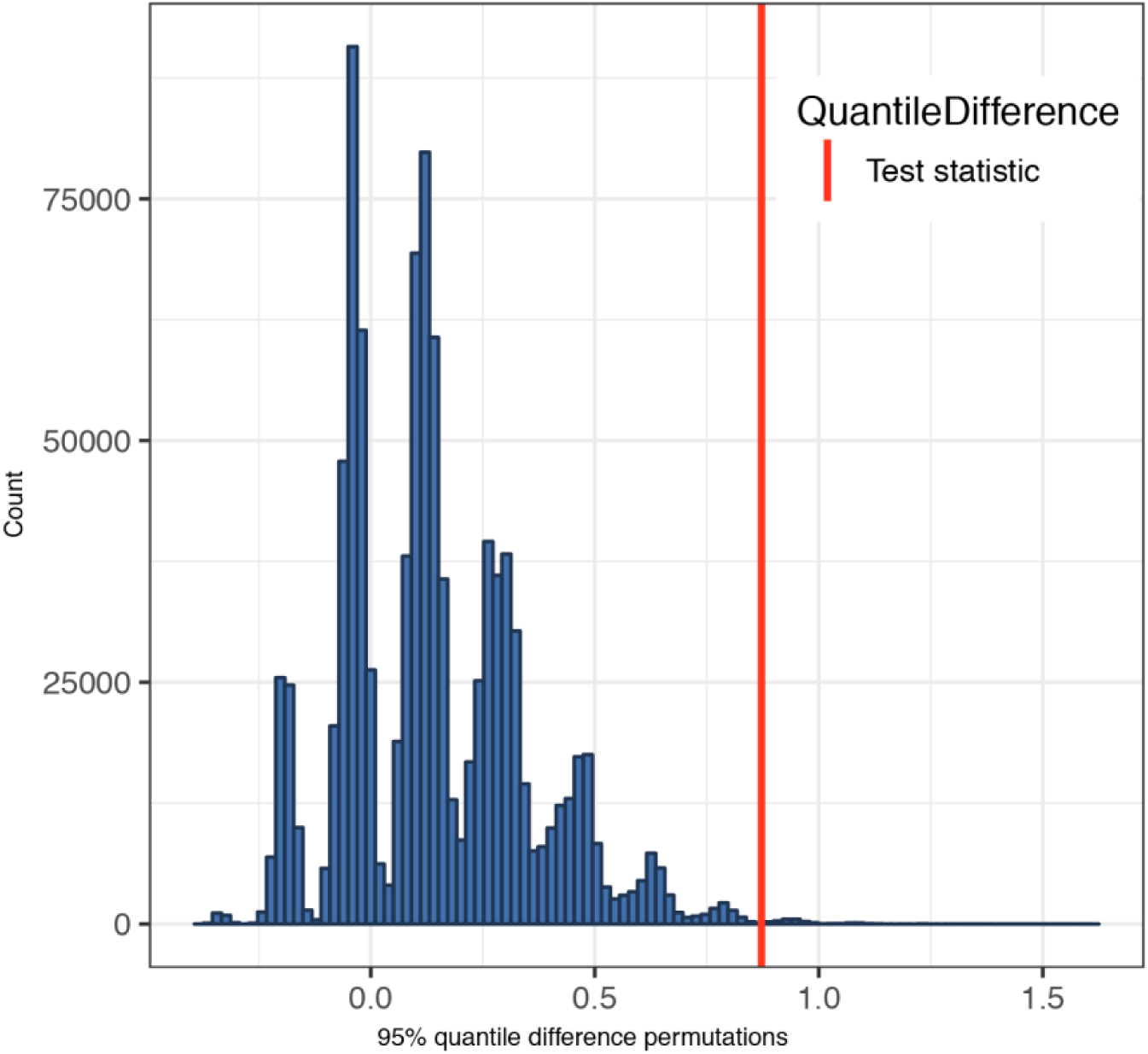
Histogram of permutations, 95% quantile difference test:
The histogram of differences between the 95% quantile of 1million permutations of the probability densities from the pairwise distances of plasmids with at least 3 antibiotic resistance genes and those with None from Figure 4. The red bar shows the observed difference in 95% quantiles between the groups.

**Supplementary Figure 5.**
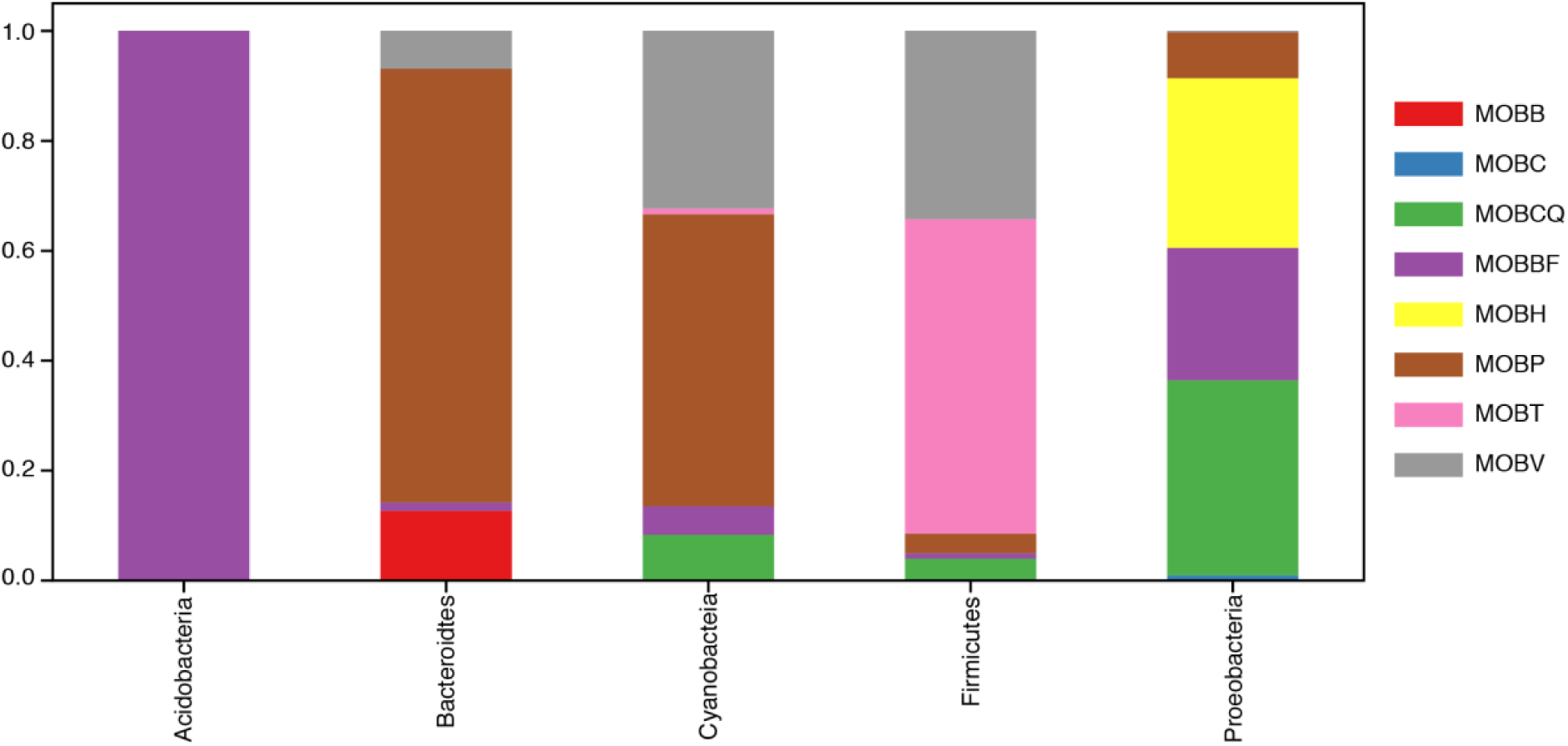
Distribution MOB types among phyla:
We show the proportion of each MOB type associated to with phyla based on a search of all known MOB types from Guglielmini et al. in the all-microbial-index.

